# Assessing Knowledge Distillation of a Multi-Emitter Localizing Neural Network for Applications in Stochastic Optical Reconstruction Microscopy

**DOI:** 10.64898/2025.12.19.694650

**Authors:** Micheal B. Reed, Reza Zadegan

## Abstract

**Background:** Super Resolution Microscopy (SRM) is a powerful method in quantitative bioscience that allows interrogation of nanoscale details. These methods require extensive imaging times on the microscope resulting in data sets on the scale of gigabytes. In order to reduce imaging times, the concentration of emitters can be increased, however that results in overlapping emitters rendering the isolation of single emitters extremely difficult. Statistical methods have been developed to deconvolute overlapping emitters, however they require parameter optimization and user expertise. Recently, Machine Learning (ML) has been developed to automate this analysis but often require larger Convolutional Neural Networks (CNN). While powerful, such models require compute and storage that would make pushing these models to compute limited devices difficult. To address this, we investigate if the dense multi-emitter localization capacity of a larger model, Deep Residual Stochastic Optical Reconstruction Microscopy (DRL-STORM), can be transferred to a smaller model, Super Resolution Convolutional Neural Network (SRCNN).

**Results:** Knowledge transfer from DRL-STORM to SRCNN did not result in an improvement of multi-emitter localization performance of SRCNN. Hint Learning (HL) was performed to facilitate knowledge transfer in a more deliberate manner. SRCNN demonstrated a limited capacity to learn an intermediate representation of the input image in the same manner as DRL-STORM and resultantly did not perform any better in its task.

**Conclusions:** Knowledge transfer was not successful between DRL-STORM and SRCNN, but evidence suggests that it is possible and may require another model besides SRCNN. A future work will investigate if hyper-parameter optimization results in greater knowledge distillation between DRL-STORM and SRCNN. SRCNN may not be ideal for multi-emitter localizations, but it can still prove effective for SRM data analysis at emitter concentrations typical of SRM experiments and is uniquely suited as a neural network model that can be deployed in compute limited settings.

## Background

Super Resolution Microscopy (SRM) empowers researchers to observe phenomena at a scale that is beyond the diffraction limit theorized by Abbe[1–4]. Stochastic Optical Reconstruction Microscopy (STORM), which relies on the random blinking of fluorescent molecules, is one implementation of SRM and has shown to be an excellent method in biological investigations[5–9]. It has been used to characterize protein distributions, subcellular component architectures, genome architectures, and many other quantitative descriptors of cellular behavior[10–13].

Because STORM requires the stochastic activation of the emitters in a noisy environment, long imaging times are often required to acquire quality data for downstream analysis[14]. One method of decreasing the acquisition time is by increasing the number of emitters in the sample[15]. Theoretically this increases the number of detected localizations which is directly related to the accuracy of localization. However, this introduces the issue of overlapping emitters which require multi-emitter fitting models such as DAOSTORM[16, 17]. Such algorithms employ the Least Squares formulation, deconvolution algorithms that use the maximum likelihood framework or Bayesian inference[18, 19]. While useful, each requires extensive data processing times in addition to sample dependent parameter optimization.

To speed up data processing and circumvent parameter tuning of these algorithms, Deep Learning (DL) models have been developed to perform single and multi-emitter localizations[20–22]. DL is a subset of Machine Learning that uses convolutional neural networks which have had enormous success in biospectroscopy, nanophotonics, and biosensing[23–26]. Deep-STORM was one of the earlier works that demonstrated that DL can indeed be used for single molecule localization[20]. DRL-STORM then aimed to address the shortcomings of Deep-STORM in live-cell applications which require dense emitter concentrations by incorporating residual connections to improve the image denoising capacity of the algorithm[27].

While successful, both models still require millions of parameters to be optimized during the training phase. This could restrict their application in situations where computational and space resources are restricted such as the current push to perform high quality, quantitative bioimaging on mobile phones and other edge devices[28–31]. Methods such as network pruning and compression have shown promise but can result in unwanted image artifacts[32–34]. Knowledge Distillation (KD) is a method where a teacher network teaches a smaller student network to perform a task with the same level of accuracy as the teacher network[35, 36]. While KD is typically performed in classification tasks, recent efforts have been made in the regression problem space[37]. Additionally, many algorithms utilize the up-sample operation to produce the super-resolved image because it allows for data imputation at the subpixel resolution level that is inaccessible at the low-resolution image[20, 22, 27]. However, this operation is computationally expensive resulting in longer processing times in addition to larger storage space requirements.

In this work, we asked two primary questions: First, we asked if DRL-STORM can serve as a teacher network to impart its multi-emitter localization capacity to Super Resolution Convolutional Neural Network (SRCNN)? SRCNN is a simple three-layer network first introduced in 2014 for image super resolution in traditional image processing. Second, we asked whether an image of finer detail compared to traditional fluorescent microscopy is still obtainable after foregoing the up-sampling procedure during the emitter localization process. The results of our study suggest that both are possible though further optimization is required.

## Methods

### A. Deep Learning

#### 1. Architecture

Two artificial neural networks were selected to assess the potential of KD from a teacher network to a student network. We selected DRL-STORM as the teacher network and SRCNN as the student network since the former has demonstrated high dense emitter localization performance while the latter’s performance is unknown[38, 39]. We also considered the relative architectural complexity between the two networks when deciding which would be the teacher. DRL-STORM contains both more parameters and diverse computational methods such as max-pooling and up-sampling compared to SRCNN. Another feature of DRL-STORM that we hypothesized would impart superior performance is the skip connection between the input and the output of the first half of the network. This skip connection allows the computed loss to be ‘seen’ by other regions of the network in addition to circumventing the ‘vanishing’ gradient issue often found in deep neural networks[40, 41]. Within the context of DL enabled SMLM, skip connections enable spatial information from the raw input image to be passed to later learned abstract features effectively grounding the learned representations in the emitter localization task[42, 43].

Comparatively, SRCNN is a simple network containing significantly fewer parameters and convolutional layers than DRL-STORM resulting in fewer stages of feature abstraction which could hinder its relative. Additionally, the structure of this network in its original intention is to perform super resolution in the traditional image processing sense where a smaller dimension image is enlarged and mapped onto a larger dimensioned image with the intention of containing more detail within a region of the high-definition image compared to its lower resolution analogue.

#### 2. Training, Validation and Test Dataset Generation

The ImageJ plug-in ThunderSTORM was used to produce five 1000×1000 images each containing emitter densities of 13 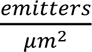, magnification of 1, and a background noise level of 200 [44]. For each of these five diffraction limited images, their super-resolved counter parts were also generated resulting in five X,Y pairs. From each of these images, 200 random 200×200 patches were obtained and were randomly assigned to the training, validation or test dataset according to a random draw which resulted in roughly an 80%:10%:10% split respectively resulting in an overall dataset containing 20,000 X,Y image pairs (Table S1). Networks were also trained with an emitter density dataset of 5 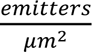 using 10,000 X,Y image pairs (Table S2).

#### 3. Training, Validation, and Testing of Models

Before the training run, the diffraction limited images were normalized using the mean and standard deviation of the entire data set. We used a similar L1L2 loss function found in previous works, however we divide the lambda parameter by 2 to remain consistent with Kim et al whose loss function informed Deep-STORM[12]. The analytical form is as follows:

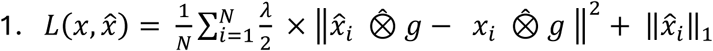

Where *x̂* is the networks prediction, *x* is the ground truth, ⨶ is a convolution with a 3×3 Gaussian kernel with a standard deviation of 1, and λ is a hyperparameter that controls the sparsity of the output. The network was implemented using the PyTorch framework. Each network training run consisted of 50 epochs with a batch size of 5. We used the Adam optimizer with a learning rate of .0001 and a scheduler which reduces the learning rate by an order of magnitude after no observable change in the loss after 5 epochs. All training and validation were performed on the gpu partition of the Longleaf computing resource hosted at UNC Chapel Hill which utilizes NVIDIA GeForce GTX 1080 with 8GB memory.

All inference tasks were performed on the ‘general’ partition of the Longleaf super-computer with the following hardware specifications: 24 x Intel(R) Xeon(R) CPU E5-2643 v3 @ 3.40GHz.

Normalized Mean Square Error (NMSE) and Structural Similarity Index Measure (SSIM) were the two metrics used to assess the performance of trained networks on the test dataset. The NMSE is defined as 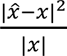 where *x̂* and x are the predicted and ground truth super resolved image respectively.

#### 4. Knowledge Distillation

Two strategies of KD were employed to investigate the efficacy of transferring knowledge from DRL-STORM to SRCNN. First, we use the Attentive Imitation Loss (AIL) which has the following form[37].

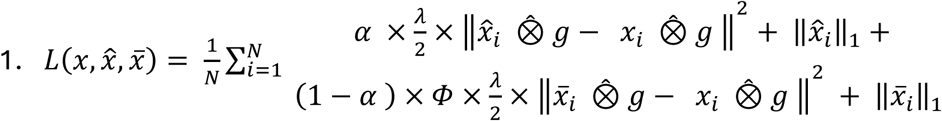

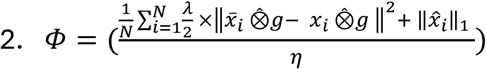

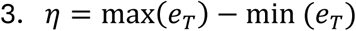

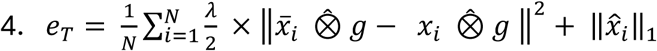

Equation 1 is used as the loss function during the student networks training. *x̂*_*i*_ is the student network’s prediction, *x̅_i_* is the teacher network’s prediction, *x*_*i*_is the ground truth, and *e*_*T*_is the vector containing the L1L2 loss values generated from the training phase of the teacher network. The alpha value in Equation 1 controls the attention of the student model. The lower the alpha value, the less emphasis the student model places on its difference between it and the ground truth. This coerces the student model to minimize the difference between its output and the teacher output. Resultantly, an alpha value of 0.1 coerces the student model to pay more attention to the teacher model, while an alpha value of 0.9 is more attentive to the ground truth

The second approach, and optionally complimentary approach, is to perform hint learning before training the student model. Hint learning is when the teacher model, which is assumedly more accurate than the student model, “teaches” the student model to have an intermediate representation that more closely resembles the teacher network. The idea is that if the student network can learn how to represent the input in a way that mirrors the teacher network, then it will be easier for the student to generate a similar final output. For this to happen, the intermediate representations of the networks must have the same dimensions. In our work, we chose the penultimate layer which contains 32 channels in both models. We hypothesized that this intermediate representation contains the de-noised information needed in order to localize the dense emitters.

We accomplished hint learning by implementing both DRL-STORM and SRCNN in two parts: from beginning to hint layer, and then the final reconstruction layer. We pursued the typical training process on this factorized version of DRL-STORM with the same loss function parameters we found to establish our baseline model DRL-STORM model. We considered the 32 channel outputs of this model to be the “ground truth” by which the SRCNN model was trained. The loss function for Hint Learning is the Mean Squared Error which is as follows:

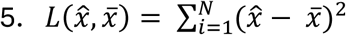

Where *x̂* is the student prediction and *x̅* is the teacher prediction. Once the hint was learned, it was put back into series with the final reconstruction layer where only the parameters of the final reconstruction layer were passed to the optimizer so that the weights through the hint layer remained static. During this final training process Equation 1 was used to optimize the weights of the reconstruction layer.

#### 5. Hyperparameter Search

To find appropriate values for the lambda hyperparameter in the L1L2 loss function and alpha in the AIL, we ran training, validation and testing experiments across a range of values sensible for both parameters. For the L1L2 loss function we investigated lambda values of 10,50,100,200,400, and 800. In regard to the alpha parameter in AIL we investigated values of 0.1-0.8 in increasing 0.1 increments.

## Results

### 1. Attentive Imitation Loss

It was imperative that we first establish a well-performing teacher model before moving forward with the KD experiments. Additionally, we wanted to ensure that the student model did not have the same performance as the teacher model so that it would have room to improve its performance. To achieve this aim, we ran a λ parameter sweep experiment where we trained the model using the different values stated in the methods section and assessed the performance of the resulting models. Figure S2 contains a box-and-whisker plot of the test metrics of DRL-STORM trained using different λ values. According to Figure S2A, a λ value of 5 results in a NMSE of ∼27%, However, increasing this value to being >25 results in a drop in NMSE to <5% indicating that that λ of 25 or greater should be used to minimize the pixel-wise difference between the predicted image and the ground truth.

To determine the optimal λ should be used, we also assessed the SSIM distributions as seen in Figure S2B. Similarly to the NMSE albeit in an inverse fashion, the SSIM score increases dramatically from ∼0.52 at λ = 5 to ∼0.75 at λ>5. Of all λ values, 100 has the highest average SSIM score of ∼0.77. Since this λ also had a low NMSE we elected to choose 100 as our lambda value across all experiments that required optimization via the L1L2 loss function.

The resulting training and validation loss curves for SRCNN and DRL-STORM using a λ=100 are shown in Figure S3. SRCNN (Figure S3A) has a higher initial error with a value of ∼6 while DRL-STORM has a lower value ∼2.5. SRCNN training and validation loss values plateaued under 10 epochs while DRL-STORM exhibited a continuous loss even though the precipitous drop was after the first epoch. Both models display loss behavior that suggested they are not overfitting as the validation loss in both models are either higher or the same value as the training loss.

Using these trained student and teacher models, we assessed their inference performance on the test data set. Figure 1 shows the comparative results of SRCNN and DRL-STORM. SRCNN displayed a lower distribution of NMSE (Fig 1A) with a mean <0.1 while DRL-STORM had a mean NMSE ∼0.3. This suggests that on a per-pixel level SRCNN performed the localization task more accurately than DRL-STORM. However, according to the SSIM, which is a more reliable metric for image reconstruction accuracy, DRL-STORM is by far the best performing algorithm with a mean SSIM score of ∼0.80 compared to ∼0.56 compared to SRCNN. We interpreted these results to suggest that DRL-STORM is the better performing model for emitter localization in dense concentration settings, which is consistent with previous reports where DRL-STORM outperformed Deep-STORM, a fellow encoder-decoder architecture[27]. Thus, we concluded that DRL-STORM and SRCNN are well positioned to be the teacher and student networks respectively.

**Figure 1:**
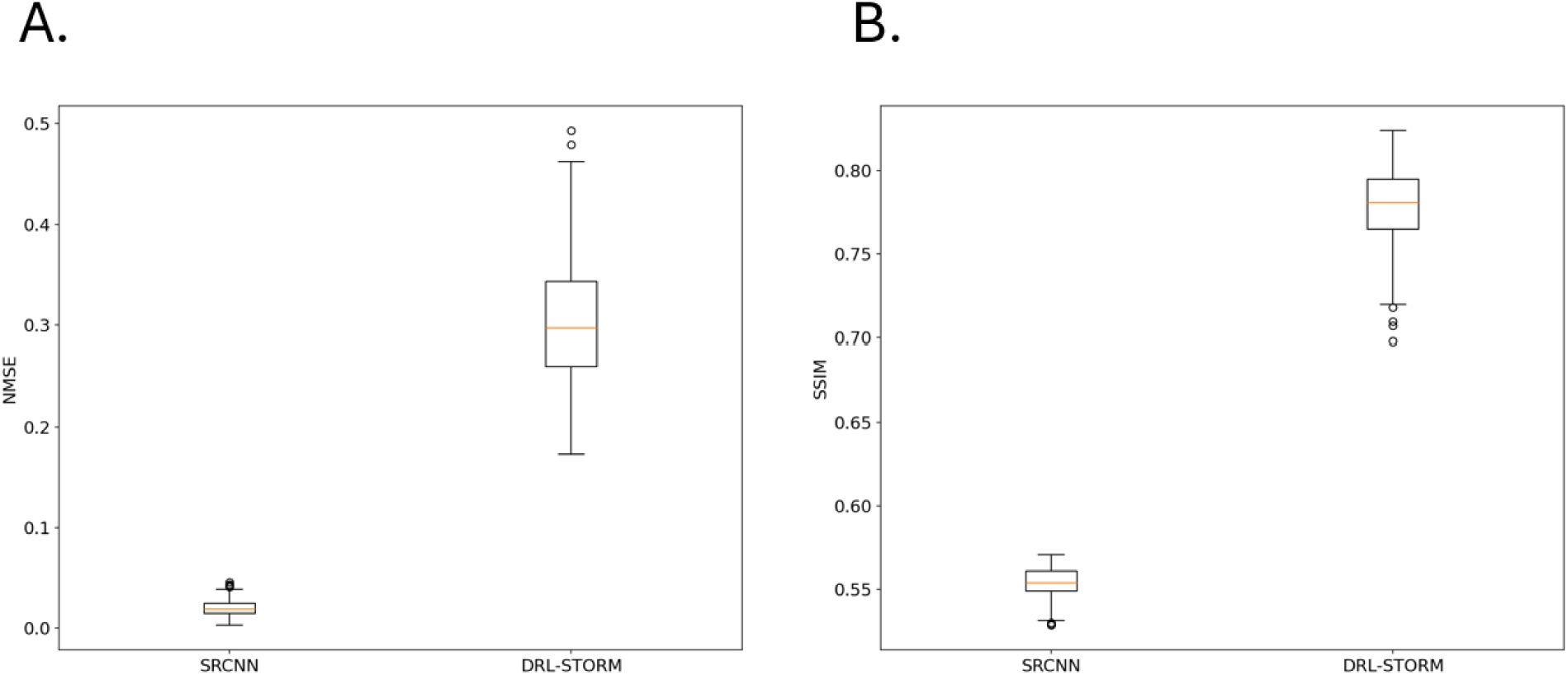
Box and whisker plots visualizing the distribution inference task metrics A) NMSE and B)SSIM for SRCNN and DRL-STORM.

Figure 2 contains the inference performance of SRCNN trained using difference alpha values in Attentive Imitation Loss (Eq1). Figure 2A shows the resultant NMSE across a range of alpha values. At lower alpha values, the NMSE is higher with an average value of ∼0.2 at 0.1 and decreases as the alpha parameter increases. The NMSE plateaus at an alpha value of 0.6 showing no marked increase in performance. Surprisingly, the mean NMSE values at alpha value’s 0.6-0.9 appear to be lower than the base SRCNN model trained only on the ground truth however the distributions do not appear to be significantly different from one another.

**Figure 2:**
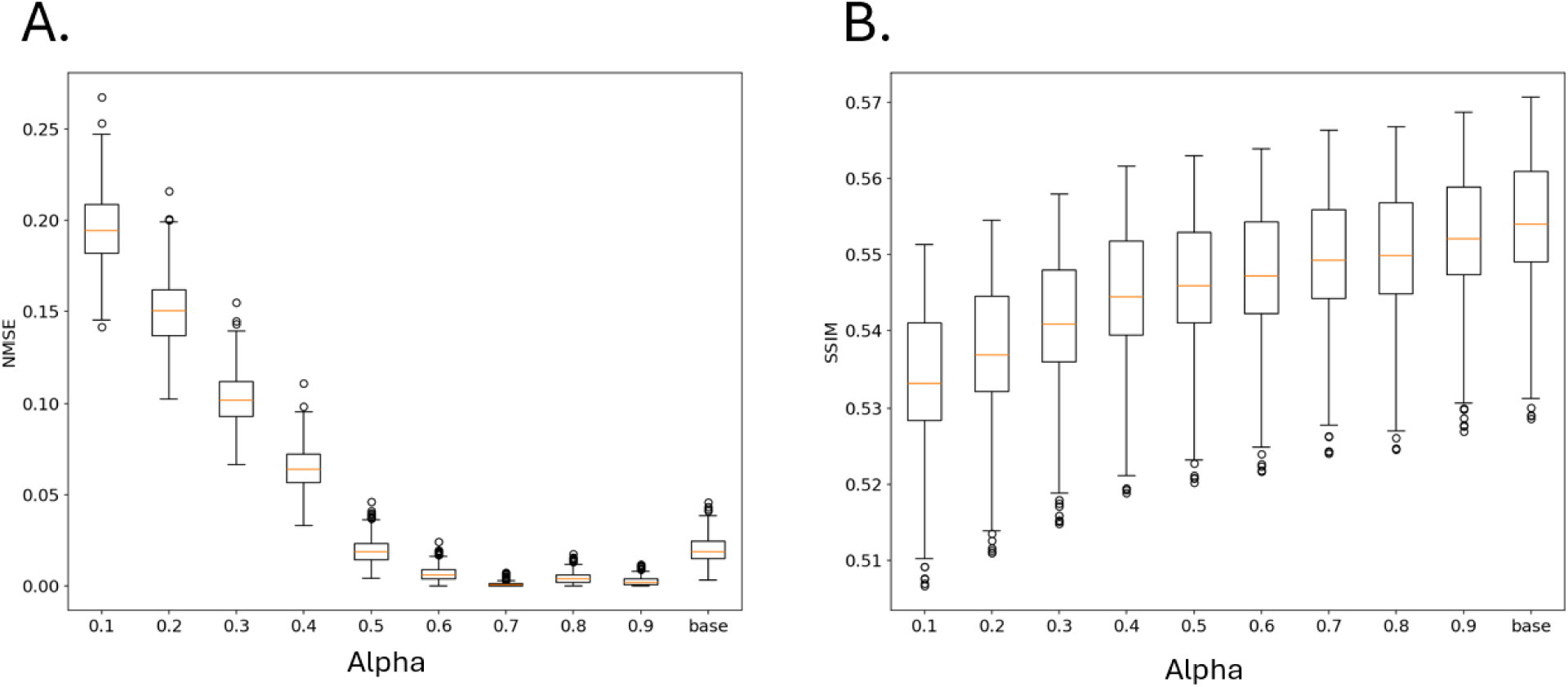
Resultant A) NMSE and B) SSIM values of SRCNN performance on the test dataset after being trained using different alpha values in the Attentive Imitation Loss function.

Figure 2B shows the relationship between the alpha value and the SSIM value of the reconstructed image of emitters. There is an observable positive relationship between SSIM and alpha values with 0.1 having an average SSIM value ∼0.53 and the baseline model having one of ∼0.55. Interestingly, the difference in inference performance between the two alpha values is negligible given that SSIM is bound between 0 and 1. We observed that the resultant SSIM distributions at alpha values between 0.7-0.9 closely resembled that of the SRCNN model trained solely on the ground truth data.

### 2. Hint Learning

Given that none of the SRCNN model outperformed the baseline model solely using AIL, we used DRL-STORM to teach SRCNN its intermediate representations, referred to as SRCNN@hint_learned in the hopes that it would lead to better inference performance (Figure S4). Figure S5 shows the training and loss validation curves generated during the hint learning experiment. There is a decrease in the difference between the two features starting with an MSE of ∼0.44 and plateauing at ∼0.38. It should be noted that this plateauing occurred in less than 10 epochs suggesting that SRCNN learned as much as it could from DRL-STORM within a short amount of training cycles.

To better understand if SRCNN@hint_learned could represent an input image in a similar fashion to DRL-STORM we used the manifold learning algorithm UMAP (Figure 3). Firstly, we observed that all clusters representing the four different assessed models were well separated. This suggested that the 32-channel representations from the different models are distinct from one another. Secondly, given that UMAP aims to preserve the global structure from high dimensional space to a lower one and, clusters that are more similar to one another are adjacent, visual inspection along the x-axis suggested that the cluster from the SRCNN@hint_learned (green) is slightly shifted more towards DRL-STORM compared to the trained SRCNN algorithm (orange). This suggested that SRCNN@hint_learned does produce a 32-channel feature that more closely resembles DRL-STORM compared to a baseline optimized SRCNN though not by much. It should be noted that both models, however, are distinct from the baseline model.

**Figure 3:**
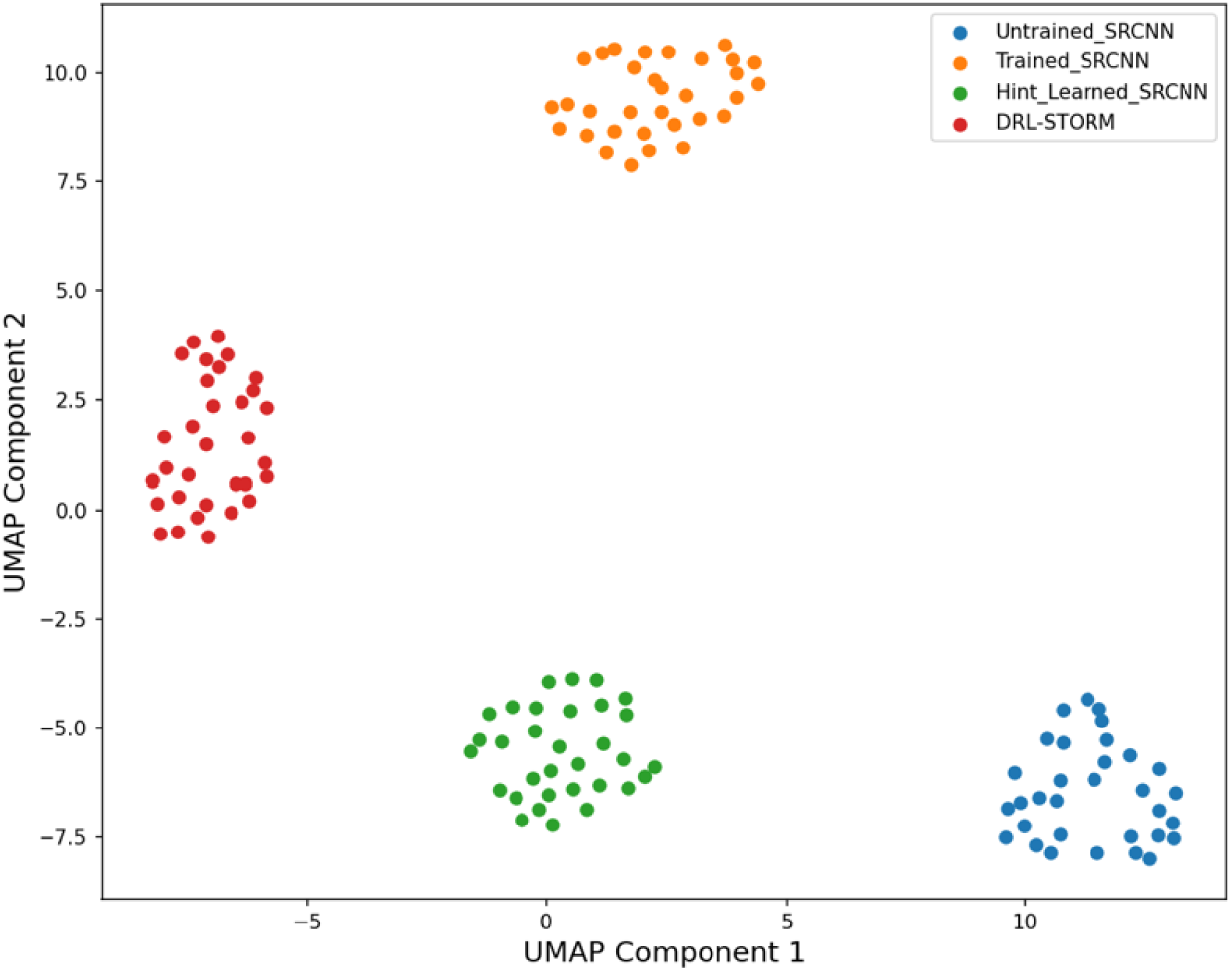
UMAP projection of the 32-channel space into a 2D space using four different models: untrained SRCNN (blue), trained SRCNN (orange), hint-learned SRCNN(green), and DRL-STORM (red)

Figure S6A shows a histogram of the pairwise distances containing the following comparison: DRL-STORM vs SRCNN@hint_learned and DRL-STORM vs SRCNN. We observed heavy overlap between the distributions though SRCNN@hint_learned was shifted to the left suggesting that the distances between it and DRL-STORM were smaller. We concluded that minimal hint learning had occurred. Next, we proceeded to train the final layer of SRCNN@hint_learned with a static network up to the hint learned layer and observed expected training and validation loss trends (Fig S7).

We then assessed the inference performance of the entire network containing the intermediate learned representation. The comparative inference behavior between the SRCNN with the learned representation and DRL-STORM mirrored that of the baseline optimized SRCNN (Fig S8). SRCNN once again had a smaller average NMSE and SSIM value compared to DRL-STORM. To investigate how respective networks would transform the diffraction limited image, we passed one image from the test dataset into both networks (Figure 4). Figure S9 show the ThunderSTORM diffraction limited image and its ThunderSTORM derived ground truth. The inferred image from SRCNN@hint_learned shown in Figure 4A can be observed to have noticeable differences from the ground truth. Primarily, there are emitters that are not well-localized and instead have a “smearing” effect across multiple pixels. This is in contrast to the DRL-STORM inferred image (Fig 4B) where the emitters are much more well localized even in the dense emitter regions. That said, a significant amount of noise is removed from both inferred images compared to the diffraction limited image (Fig S9A). Since we did not observe any marked improvement between SRCNN@hint learned and the optimized SRCNN model we proceeded with the latter for the remainder of the experiments.

**Figure 4:**
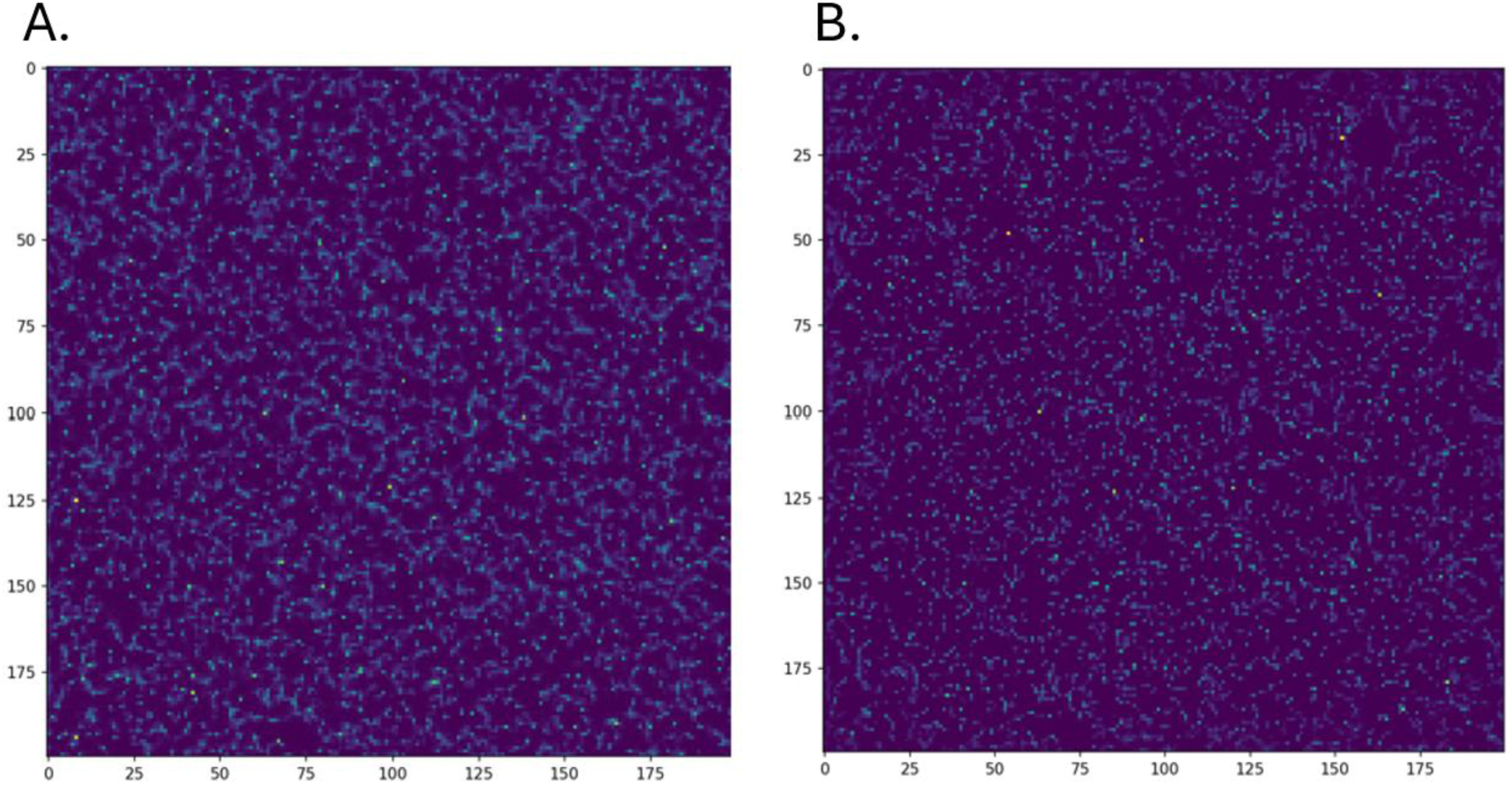
Localized emitter image produced by A) SRCNN@hint_learning and B) DRL-STORM when a diffraction limited image of emitters is passed to it.

### Image Reconstruction

### Simulated Microtubules

The trained DRL-STROM and SRCNN were then applied to perform the SRM reconstruction task on two high density emitter datasets, one simulated and the other experimentally gathered, hosted on the European Super Resolution Microscopy Hub.

Figure 5 contains the simulated diffraction limited image (Fig 5A), the ThunderSTORM (Fig5B), SRCNN (Fig 5B), and DRL-STORM (Fig 5C) super resolved images of the simulated high density microtubule data set. Compared to the ThunderSTORM image, SRCNN and DRL-STORM both contained higher amounts of noise evidenced by the haze observed in the empty space. Additionally, DRL-STORM had significant amounts of linear artifacts in the vertical and horizontal direction. While noise was present, the microtubules were properly reconstructed even capturing the distal microtube at the edge of the image that ThunderSTORM failed to detect.

**Figure 5:**
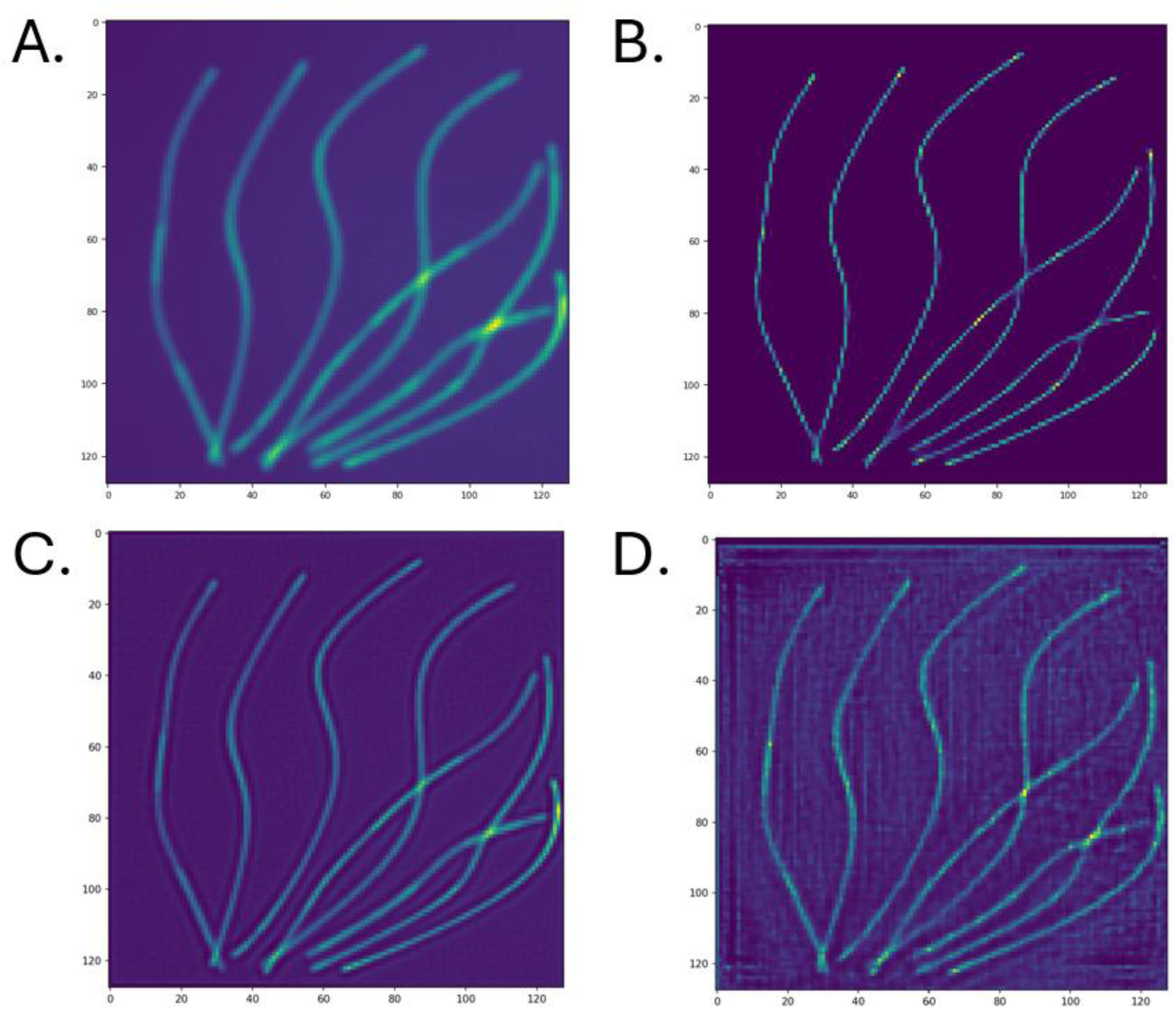
The A) diffraction limited image B) ThunderSTORM, C) SRCNN and D) DRL-STORM reconstructed super resolution image of a high-density microtubule data set. C and D were trained on datasets containing a high emitter concentration of 13 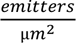

Observing the cross-profile plot, one can observe the resolution of the features in SRCNN and DRL-STORM was still a significant improvement over the diffraction limited image (Fig S11A). The widths of the peaks were narrower than the diffraction limited image but were slightly larger than the ThunderSTORM image. Evidence of the background haze in SRCNN and DRL-STORM can be seen in the cross-profile plot as they both contained higher pixel values in locations that ThunderSTORM has assigned 0’s.

Figure 6 shows the Super-Resolved image of an experimentally measured high density microtubule data set using the different models explored in this work. We observed similar patterns such as the high amount of linear and vertical artifacts, perhaps even more present than the simulated microtubule dataset (Fig 6B). We also observed artifacts in SRCNN around the perimeter of the image. Visual inspection suggested that SRCNN (Fig 6C) resulted in a clearer, more faithful representation to ThunderSTORM compared to DRL-STORM.

**Figure 6:**
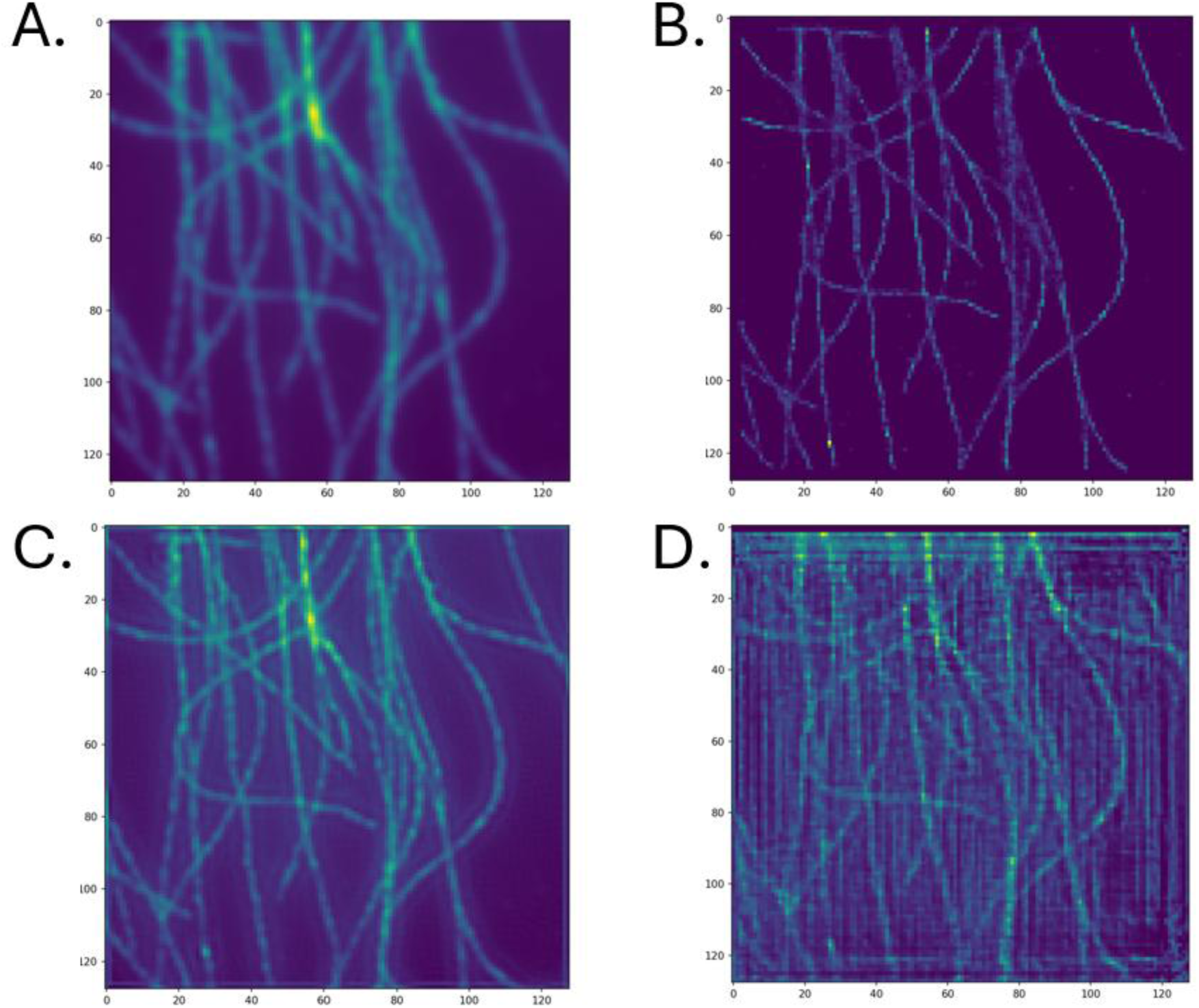
The A) diffraction limited image B) ThunderSTORM, C) SRCNN and D) DRL-STORM reconstructed super resolution image of an experimental microtubule data set. C and D were trained on datasets containing a high emitter concentration of 13 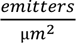

Examination of the intensity cross-profile for the experimental dataset (Fig S11B) confirmed the qualitative observation of the increased amount of noise in the image reconstruction. Like the simulated dataset, DRL-STORM and SRCNN both contained higher pixel values at locations that ThunderSTORM had set to 0. Additionally, the effect of the linear artifacts in DRL-STORM can be observed as multiple peaks are present in the plot which are not in the ThunderSTORM profile. In contrast, SRCNN did not contain these spurious peaks but followed the outline of ThunderSTORM. Both SRCNN and DRL-STORM had a higher resolution compared to the diffraction limited image evidenced by the narrower peaks compared to the diffraction limited image. Altogether, despite the high amount of noise (requiring a lowering of the lambda parameter) the performance of SRCNN and DRL-STORM appeared to achieve super resolution capacity.

Previous works have removed the edge artifacts by lowering the density of the emitters in the training data[27]. We re-trained both models using an emitter density of 5 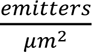 and a lambda value of 10. Then the image reconstruction task on the experimental and simulated data set was repeated. The results of the former can be seen in Figure 7. In this instance, DRL-STORM produced an image with significantly fewer artifacts compared to when it was trained with a more concentrated emitter density (Fig 7D). However, horizontal and vertical artifacts were still present in the image suggesting that its architecture had a role in imparting these artifacts. The reconstruction result from SRCNN (Fig 7C) closely resembled its result from its counterpart trained on a higher emitter density. The primary difference was that in Fig 6C, SRCNN contained high background along the contours of the microtubules whereas in Figure 7C the higher background signal was localized to the bottom right-hand corner and was less pronounced along the microtubules. The cross profile of intensity values showed that the widths of the microtubules for SRCNN and DRL-STORM were narrower compared to the diffraction limited image but also contained more noise than ThunderSTORM. Notably, SRCNN contained more noise compared to DRL-STORM which supports the claim from the original paper that the first half of the network removed the noise in the image enabling noise-free image reconstruction[27].

**Figure 7:**
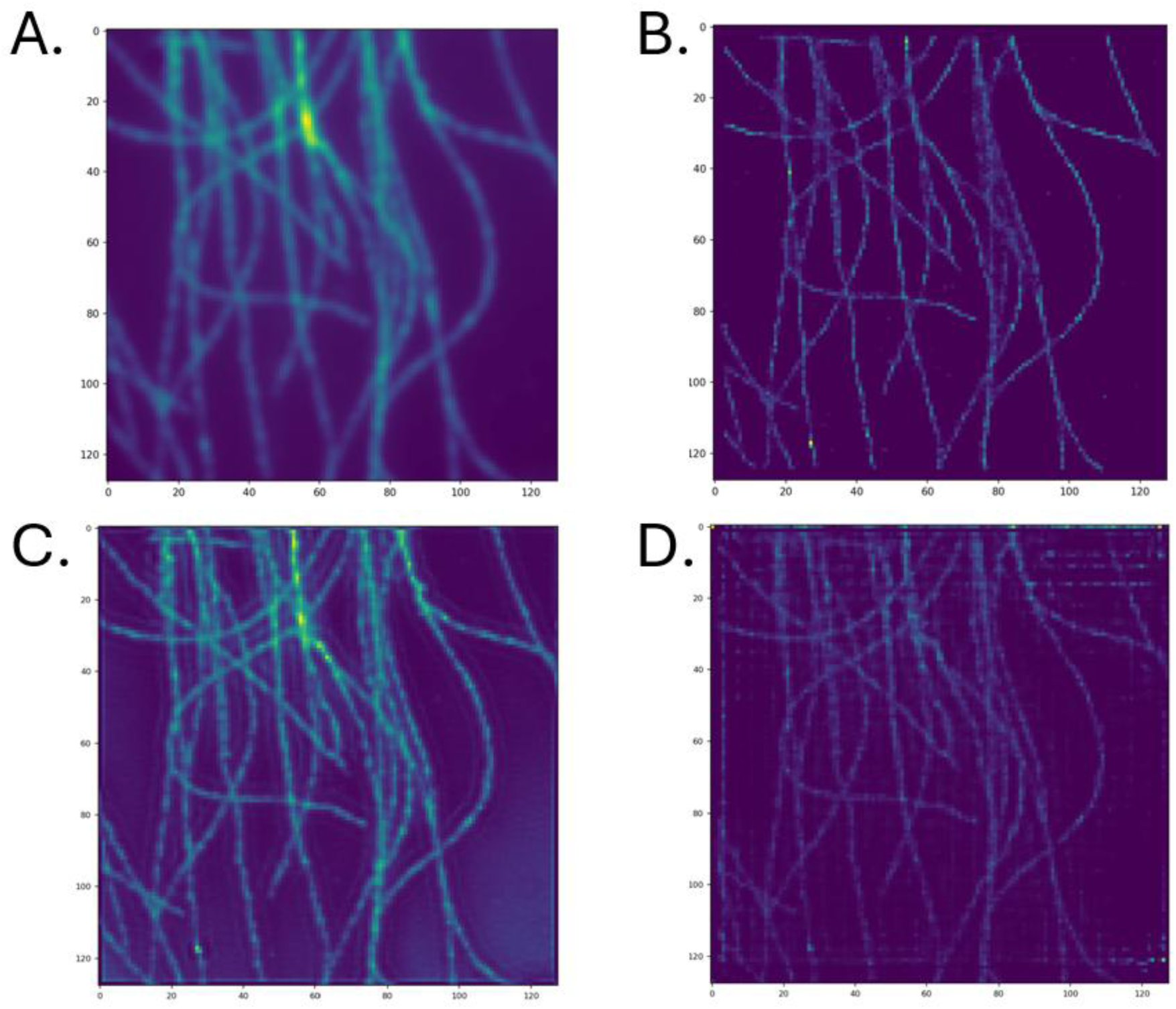
Figure 6: The A) diffraction limited image B) ThunderSTORM, C) SRCNN and D) DRL-STORM reconstructed super resolution image of an experimental microtubule data set. C and D were trained on datasets containing a high emitter concentration of 5 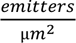

## Discussion

In this work we investigate the possibility of whether a teacher network, DRL-STORM, could teach a student network, SRCNN, how to perform the single molecule localization task with high accuracy in a dense emitter environment. Our results suggest that SRCNN did not learn how to perform this task any better than it would have if trained on the ground truth data set itself. While seemingly unsuccessful we made several findings that have implications

Firstly, SRCNN reached its peak emitter performance during the knowledge distillation process using an alpha value of 0.6 in the AIL. This suggests that SRCNN does not need access solely to the ground truth data set. Additionally, even at low alpha values SRCNN has an emitter localization performance that is comparable to its optimized form. Together, these results suggest that SRCNN could hypothetically be trained solely by an optimized teacher network and not the ground truth data set. This could have implications in situations where data storage capacity is limited and cannot hold entire data sets.

While SRCNN displayed a minor capacity to learn the intermediate representation that DRL-STORM was using (Fig 3), it seemingly was not enough to improve the performance of the model. We reason that SRCNN was unable to learn the intermediate representation because of its architectural simplicity. It should be noted that the 32-channel representation is preceded only by one layer, a 64-channel representation. Additionally, SRCNN performs a 1×1 convolution in this 64-channel representation to produce the 32-channel representation. This differs from DRL-STORM where the 64-channel representation undergoes a convolution operation and a 2×2 up sampling.

It may be the case that preceding layers in both respective models are too different from one another and resultantly the weights in the 32-channel representation are coerced into spaces that are too different. One potential avenue to ameliorating this issue is to use a more complex student network that still has less parameters than DRL-STORM. One candidate network is Deep-STORM. Like DRL-STORM, Deep-STORM is an encoder-decoder architecture and so its structure more naturally aligns with DRL-STORM.

Another method that may facilitate knowledge transfer is to perform the intermediate representation learning between the teacher and the student network at multiple points in the student work instead of just at one point as explored in this work. That way the student network experiences more of a guided learning at the behest of the teacher instead of relying on the student to learn one representation. Consequently, this will require multiple instances of knowledge transfer. Additionally, there is also a number of hyperparameters that could be optimized, such as the learning rate, during the intermediate learning process.

This work also sheds light on the relationship between the lambda parameter in the loss function and the quality of the reconstruction that we have not yet seen in other works. The higher the lambda parameter the more the signal from the original diffraction image was maintained in the final reconstruction. One should be careful when interpreting SSIM and NMSE values obtained when using higher lambda values. We presented results using a higher lambda value that resulted in emitter localization metric scores that would lead an investigator to assess that the model is performing accurately. However, the resulting image reconstructions were dissatisfactory as they were riddled with artifacts. Earlier works have observed the same issue and have either trained their data on less concentrated emitter densities or removed emitters from the boundaries. Both methods suggest that the presence of emitters on the borders of the images is responsible for the artifacts in the reconstructed image. We suggest that investigators practice caution when interpreting the test metrics and instead look at the image reconstruction result as an authoritative parameter on how to optimize the lambda parameter. One direction would be to subject the lambda parameter value to an optimization algorithm, such as gradient descent or ascent, according to the resulting SSIM value between the model’s inferred image and the ground truth.

While the reconstructed images of DRL-STORM and SRCNN did not result in the same image quality as ThunderSTORM they contained finer detail resolution compared to the simulated fluorescent microscope image. This improved resolution was even more pronounced when trained on a lower emitter density before image reconstruction. This suggests that image reconstruction that rivals the quality of the majority of super resolution pipelines is not possible without the up-sampling operation.

The reconstructed images of SRCNN and DRL-STORM both contained residual noise in the case of the former or image artifacts for the latter. We hypothesize this is a consequence of the architectural differences. SRCNN does not employ residual connections that can assist with denoising the diffraction limited image. This could result in maintaining noise from the original image in the emitter localized images. DRL-STORM contains down-sampling and up-sampling operations which are typically found in CNN’s. However, these operations can result in artifacts when signal is located along the borders of the image. One way to clean both reconstructed images is to generate one-shot noise removal models such as Noise2Void or any other model that aims to accomplish the same task[45–48].

One potentially interesting research direction would be to investigate if SRCNN can perform the regression task of solely predicting the x,y localizations of the emitters instead of inferring the image itself. This task has been demonstrated in a recent work where ResNet has been trained to predict the x,y locations of emitters in a diffraction limited spot[21]. While related, this is a fundamentally different problem as it is predicting a N x 2 matrix, if solely restricted to the xy pairs, as opposed to a N x N pixel image. In that way, the up-sampling operation is not necessary and instead, the output table would be used to populate a single image with spikes at the predicted positions followed by a gaussian convolution to account for the point spread function of the microscope.

## Conclusions

SRCNN was not able to perform emitter localization in dense emitter settings as accurately as DRL-STORM. Even after knowledge distillation between DRL-STORM and SRCNN, SRCNN’s performance did not rival that of DRL-STORM. Hint Learning was used to facilitate KD in a more deliberate fashion and while minor knowledge transfer was observed ultimately the performance still was not improved. However, the observed trend suggests that it is possible. Future experiments would explore other architectures aside from SRCNN to assess the possibility of knowledge transfer.

Nonetheless we hypothesize that the best case uses for SRCNN are when the emitter density is comparably low, in the order of typical SMLM experiments. While this will not decrease the imaging time of these experiments, its relatively small size makes it a prime candidate for a workhorse SMLM algorithm on microscopes engineered from edge devices. We envision additive manufacturing methods such as 3D printing, microelectronics, network protocols, and low-footprint ML models such as SRCNN can be integrated into a smart robotic microscope that can be an essential tool in the analyst toolkit for in-field investigations[49–56].

## Lists of abbreviations

DL: Deep Learning
ML: Machine Learning
SRM: Super Resolution Microscopy
SRCNN: Super Resolution Convolutional Neural Network
DRL-STORM: Deep Residual Stochastic Optical Reconstruction Microscopy
SMLM: Single Molecule Localization Microscopy
NMSE: Normalized Mean Squared Error
SSIM: Structural Similarity Index Measure
KD: Knowledge Distillation
UMAP: Uniform Manifold Approximation and Projection
STORM: Stochastic Optical Reconstruction Microscopy

## Declarations

This work is supported by NIH award # 1R16GM145671 and NSF award # MCB 2027738. This work was performed at the Joint School of Nanoscience and Nanoengineering (JSNN), a member of the National Nanotechnology Coordinated Infrastructure (NNCI), which is supported by the National Science Foundation (Grant ECCS-2025462).

## Author information

### Contributions

Conceptualization: MBR, RZ; data generation, collection and analysis: MBR; methodology: MBR, RZ; funding acquisition: RZ, software development: MBR, RZ, writing original draft: MBR; supervision: RZ

## Ethics declarations

Authors declare no competing interests.

