## Supplementary Figures for "Assessing Knowledge Distillation of a Multi-Emitter Localizing Neural Network for Applications in Stochastic Optical Reconstruction Microscopy"

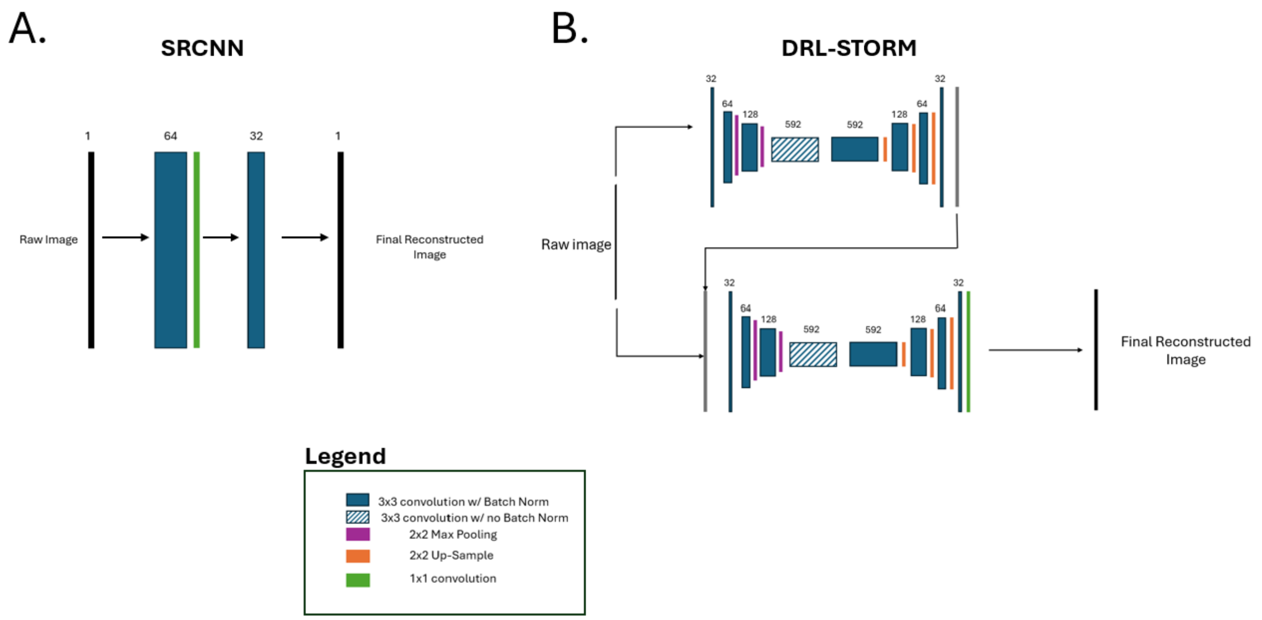


Supplementary Figure 1: Architecture of A) Super Resolution Convolutional Neural Network (SRCNN) and B) Deep Residual Learning Stochastic Optical Reconstruction Microscopy (DRL-STORM)

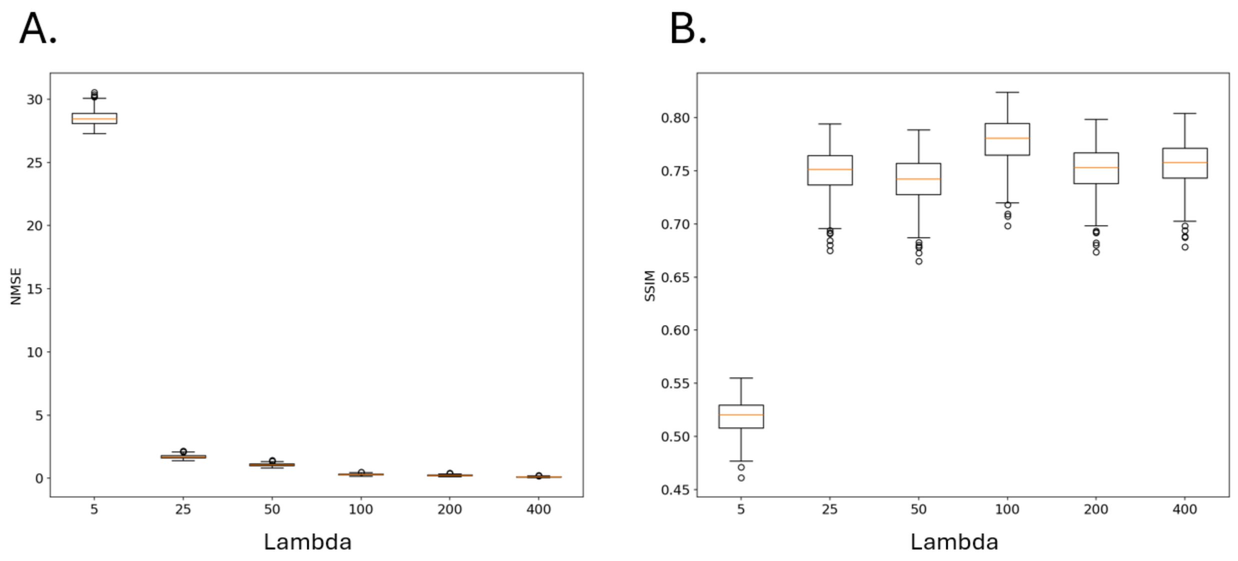


Supplementary Figure 2: Box and Whisker plots visualizing the performance of DRL-STORM on the test data set trained using different lambda hyperparameters for the L1L2 loss function. Two metrics are used to assess the inference performance: A) Normalized Mean Square Error (NMSE) and B) Structural Similarity Measure (SSIM).

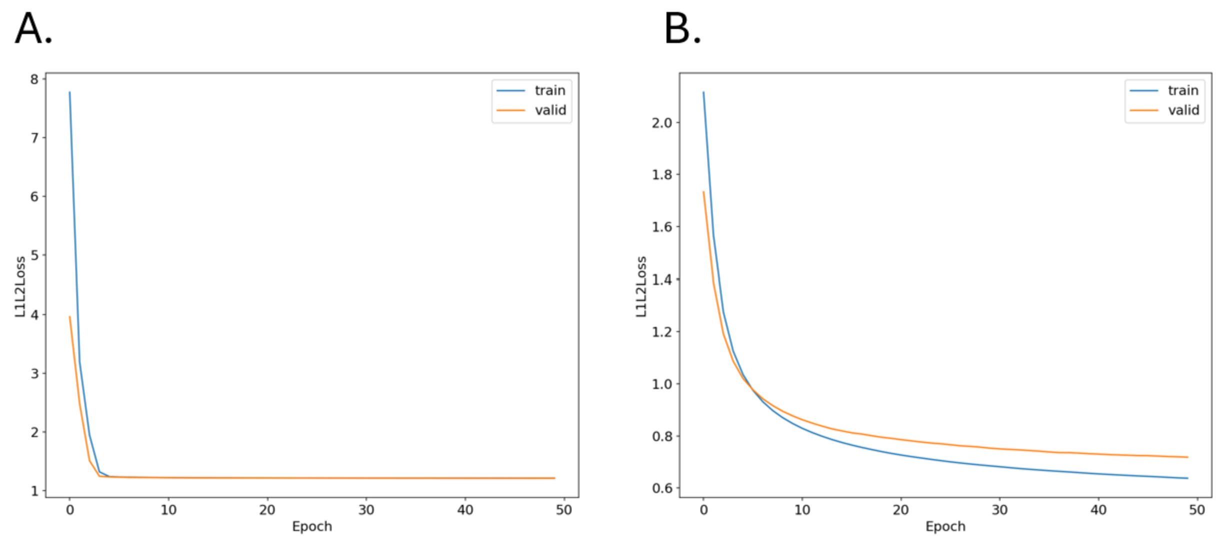


Supplementary Figure 3: Training and Validation loss curves for A) SRCNN and B) Deep Residual Learning Stochastic Optical Reconstruction Microscopy (DRL-STORM)

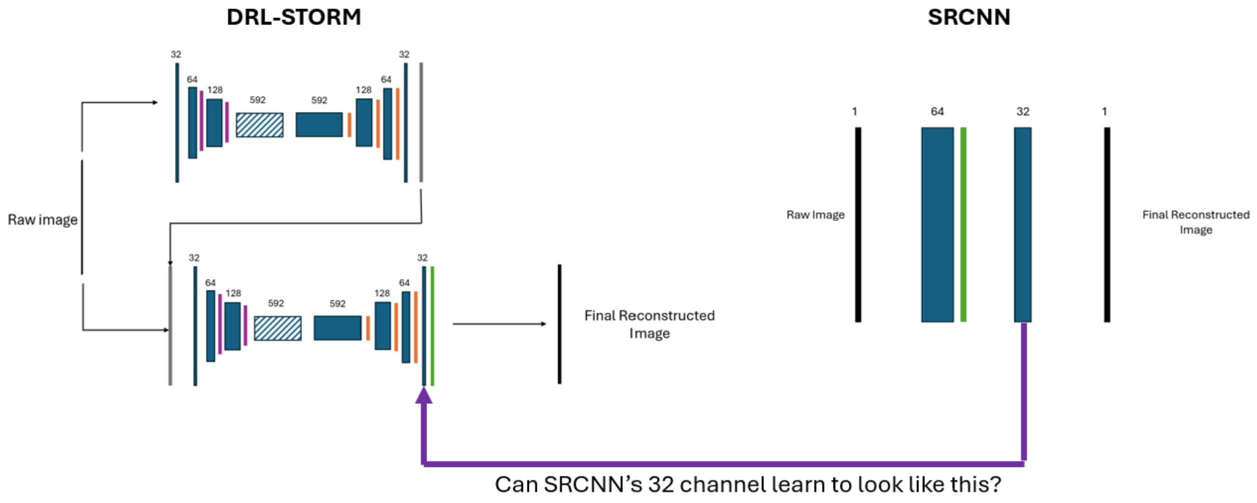


Supplementary Figure 4: Schematic showing which SRCNN intermediate representation is trying to learn from DRL-STORM.

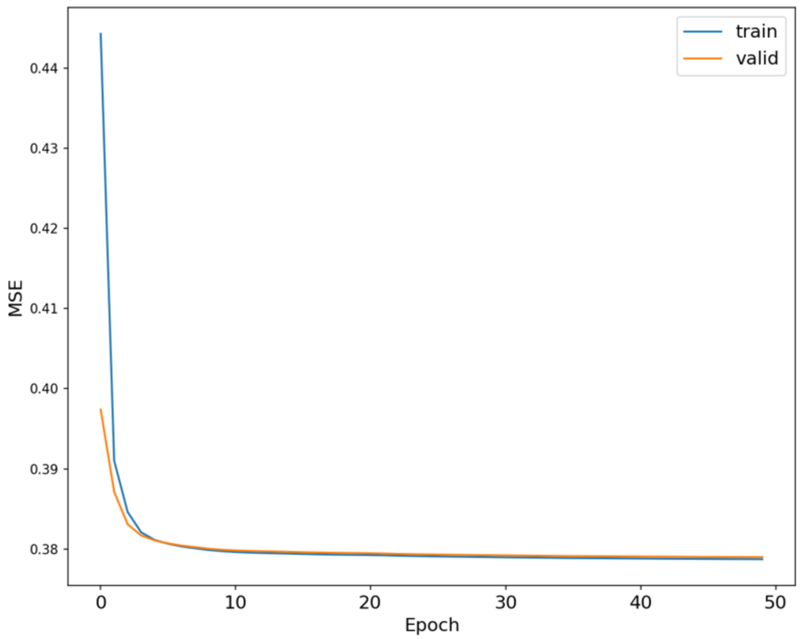


Supplementary Figure 5: Training and Loss curves with MSE as the metric between SRCNN's and SRL-STORM's 32-channel features.

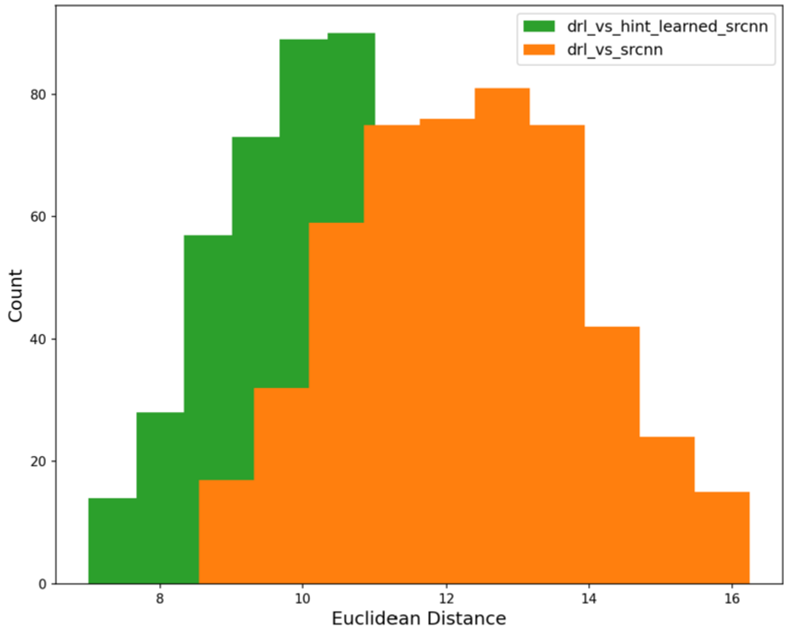


Supplementary Figure 6: Histograms of the pairwise Euclidean distances between the 32 channel intermediate representation between DRL-STORM and either base SRCNN or SRCNN@HintLearning.

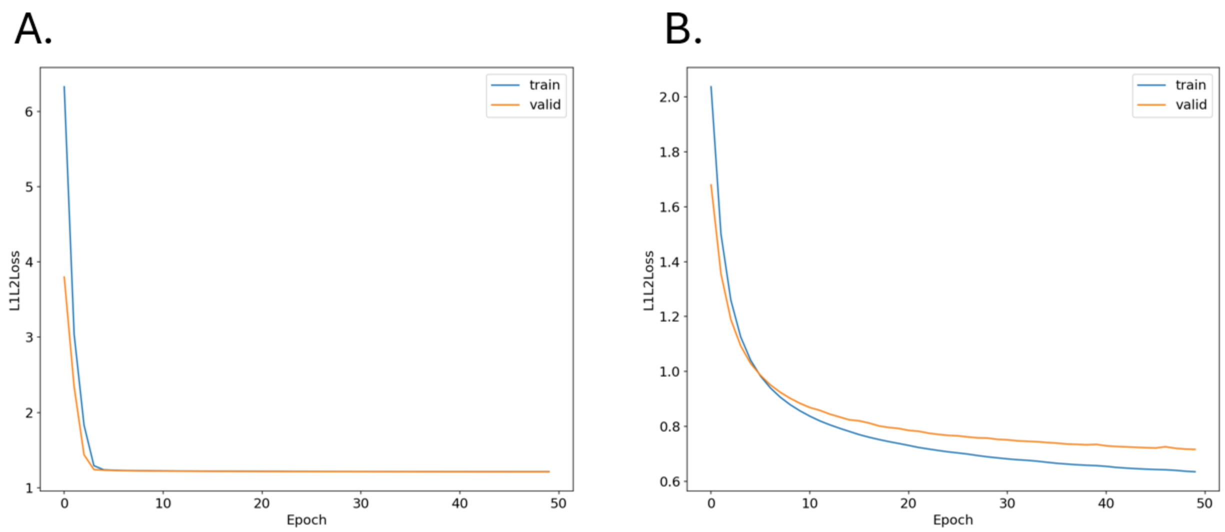


Supplementary Figure 7: Training and Loss curves of the reconstruction later for A) SRCNN@HintLearning and B) DRL-STORM.

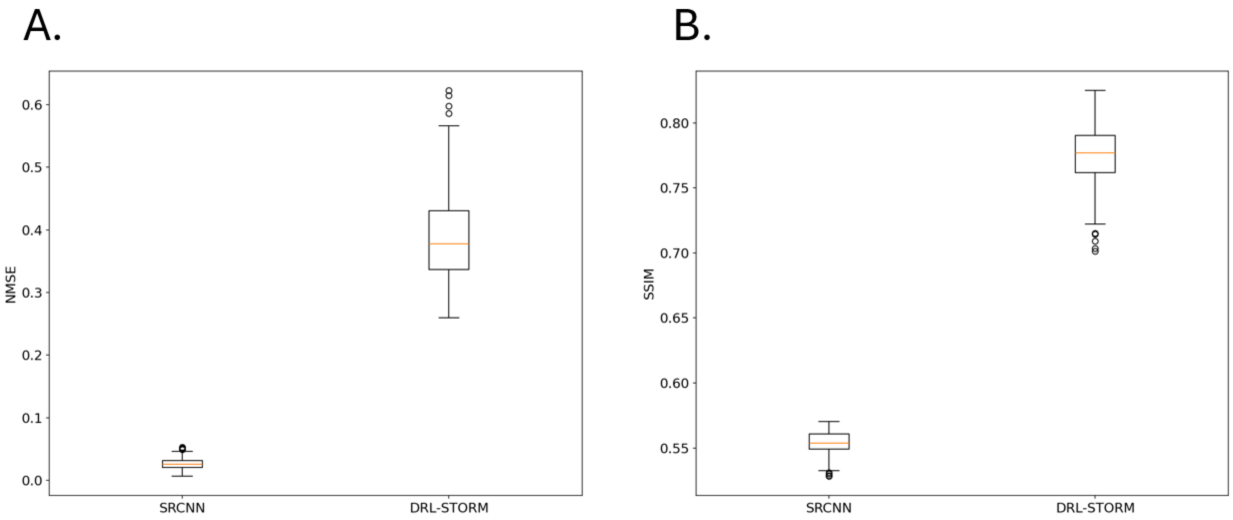


Supplementary Figure 8: Box and whisker plots visualizing the distribution inference task metrics A) NMSE and B)SSIM for SRCNN w/ learned intermediate representation and DRL-STORM.

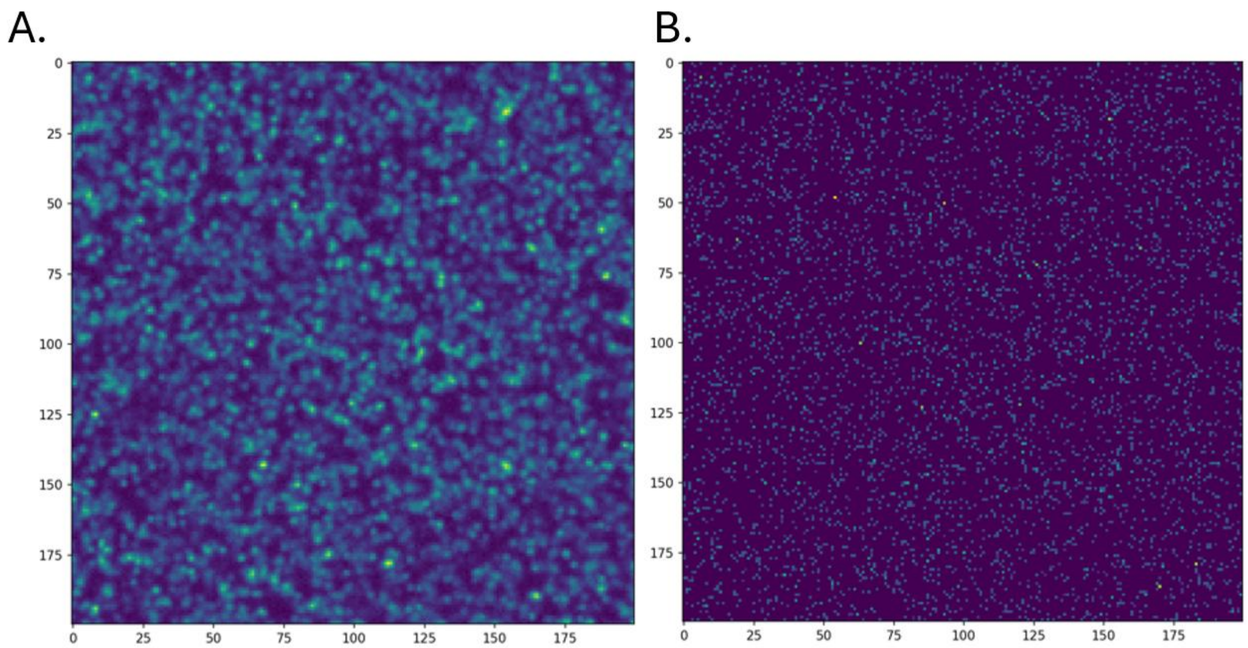

Supplementary Figure 9: Representative A) X and B)Y pair of the dense emitter localization data from ThunderSTORM.

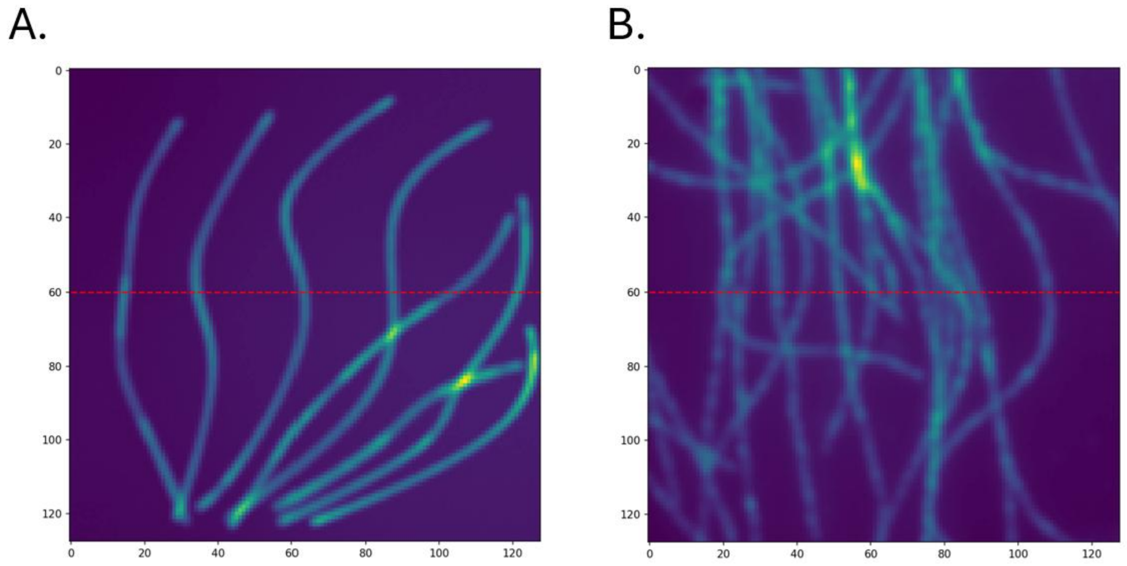

Supplementary Figure 10: A) Simulated and B) Experimental images containing a red hashed line containing the location of the extracted cross profile at pixel 60.

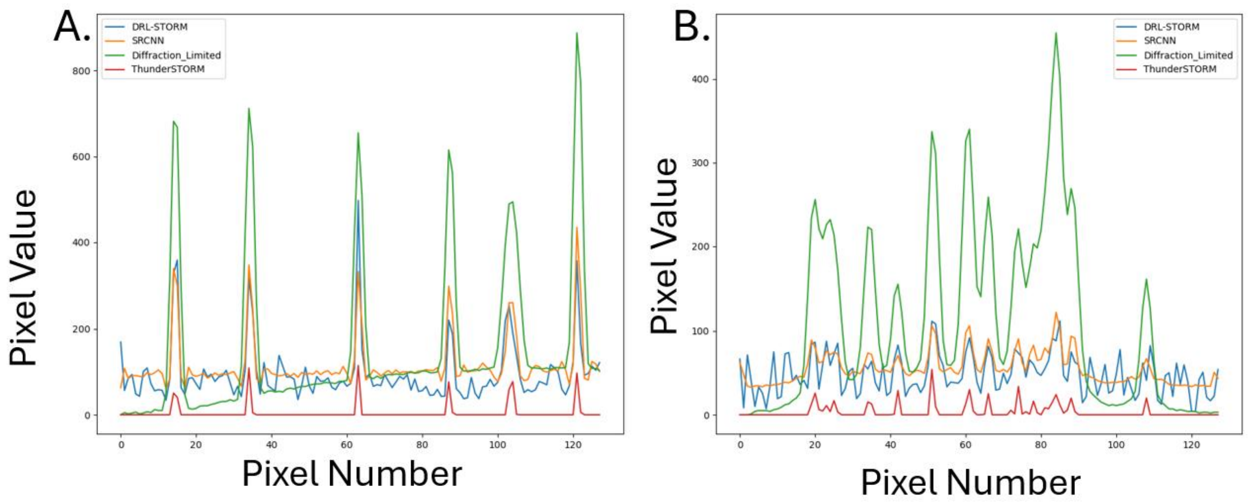


Supplementary Figure 11: Cross profile of intensity values at pixel 60 for the simulated diffraction limited image (Fig S10A and B) and the reconstructed image using ThunderSTORM, DRL-STORM, and SRCNN. SRCNN and DRL-STORM were trained using an emitter density of $13\frac{emitters}{\mu m^2}$ with a background noise of 200. The profiles are for the A) simulated and B) experimental dataset.

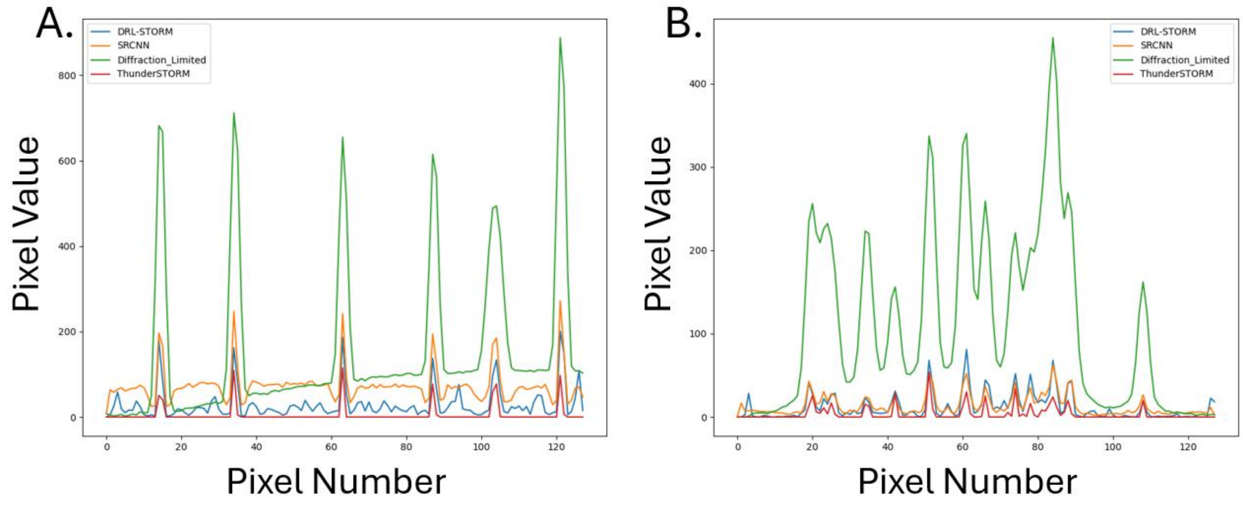


Supplementary Figure 12: Cross profile of intensity values at pixel 60 for the simulated diffraction limited image (Fig S10A and B) and the reconstructed image using ThunderSTORM, DRL-STORM, and SRCNN. SRCNN and DRL-STORM were trained using an emitter density of $5\frac{emitters}{\mu m^2}$ with a background noise of 100. . The profiles are for the A) simulated and B) experimental dataset.

Table S1 Data Partitioning for 13 $\frac{emitters}{\mu m^2}$

| Dataset | Number of Images |
|---|---|
| Training | 15,965 |
| Validation | 2073 |
| Testing | 1962 |

Table S2 Data Partitioning for 5 $\frac{emitters}{\mu m^2}$

| Dataset | Number of Images |
|---|---|
| Training | 8,037 |
| Validation | 964 |
| Testing | 999 |
